# Exploiting heterogeneity in coupled, two plasmid systems for dynamic population adaptation

**DOI:** 10.1101/2023.02.19.529072

**Authors:** Shalni Kumar, Andrew Lezia, Jeff Hasty

## Abstract

In synthetic multi-plasmid systems, it is standard to use only plasmids with orthogonal replication mechanisms to avoid phenotypic heterogeneity and ensure plasmid stability. In nature, however, microbial populations actively exploit heterogeneity to survive in fluctuating environments. Here we show that the intentional use of distinct plasmids with identical origins of replication (oris) can help an engineered bacterial population adapt to its environment. We find that copy number coupling between distinct plasmids in such systems allows for copy number buffering of an essential, but high-burden construct through the action of a stably maintained, nonessential plasmid. Plasmid coupling also generates population state memory without additional layers of regulatory control. This work reimagines how we design synthetic populations to survive and adapt by strategically giving control back to the cells.

---

Heterogeneous gene expression in cellular populations is typically viewed as a challenge to synthetic biologists striving for tight control of cellular phenotypes. While population variation can be an obstacle to engineering biology, it is pervasive in natural systems^1^. Whether driven by stochastic gene expression differences or environmental stress, living systems often strategically rely on heterogeneity to improve population survival via division of labor or bet-hedging. For example, in some biofilm forming species, more motile individuals focus on population expansion while others contribute to biofilm maintenance^2^. Additionally, some microbial populations utilize bet-hedging, a strategy where a sub-population of cells are more suited to a potential future environment at the cost of fitness in their current surroundings^3, 4^. Bet-hedging is commonly associated with spore-formation, carbon metabolism, and antibiotic persistence in bacterial populations^5^. Regardless of the mechanism, a fundamental benefit of population heterogeneity is increased adaptability to fluctuating environmental conditions^6^. The widespread use of heterogeneity in nature suggests that synthetic genetic circuits could exploit gene expression noise to adapt to different environmental conditions and carry out complex functions^7, 8^.

One source of noise commonly encountered in synthetic biology is variation in plasmid copy number (CN). Until recently, plasmid CN was an overlooked regulation strategy for tuning genetic circuits. Now researchers have developed multiple ways to use plasmid CN for genetic circuits, including creating strain libraries that exhibit different CNs^9^, designing plasmids whose CN can be tuned with small molecule inducers^10–12^, and using engineered DNA cutting to dynamically change plasmid CN over time^13^. In this work, we intentionally engineer plasmid CN noise as a tool for synthetic gene circuits by exploiting overlapping plasmid replication mechanisms.

Plasmid CN is commonly controlled by built-in negative feedback loops that act to inhibit plasmid replication when CN exceeds some set point^14^. For instance, the widely used ColE1-origin of replication (ori) makes use of an antisense RNA that inhibits replication by selectively binding an RNA pre-primer that is essential for replication^15^. Since plasmids with the same ori share the same regulation strategy, they cannot be distinguished during replication, which can increase CN heterogeneity and facilitate loss of one plasmid. Plasmids that share replication mechanisms are referred to as incompatible because they cannot be stably maintained by a cell in the absence of external selection. Although it is widely accepted that plasmids with the same ori are unstable, some research suggests that plasmids from the same incompatibility group can persist for extended periods of time without selection? and some commonly used to measure plasmid loss rate may have overestimated plasmid loss rates^16^.

In this work, we investigate the ability of two plasmid systems with shared oris to create population heterogeneity and enable adaptation to different environmental conditions (Fig.1). We show that populations with heterogeneous gene expression created by plasmid ori redundancy rapidly adapt to stressors such as antibiotics and engineered cell lysis. We also show that in these shared ori systems a non-essential plasmid can act as a tuning knob for the CN of an essential plasmid, a concept we name *plasmid buffering*. In certain environments, we find that plasmid buffering leads to increased plasmid stability for incompatible plasmids compared to compatible ones, contradicting conventional wisdom that plasmid incompatibility always leads to faster plasmid loss. We also explore the dynamics of CN adaptation, specifically looking at how changes in CN persist following a transient selection. Overall, this work shows how plasmid incompatibility can create heterogeneity in a bacterial population, enabling copy number adaptation to different environments.

**Figure 1:**
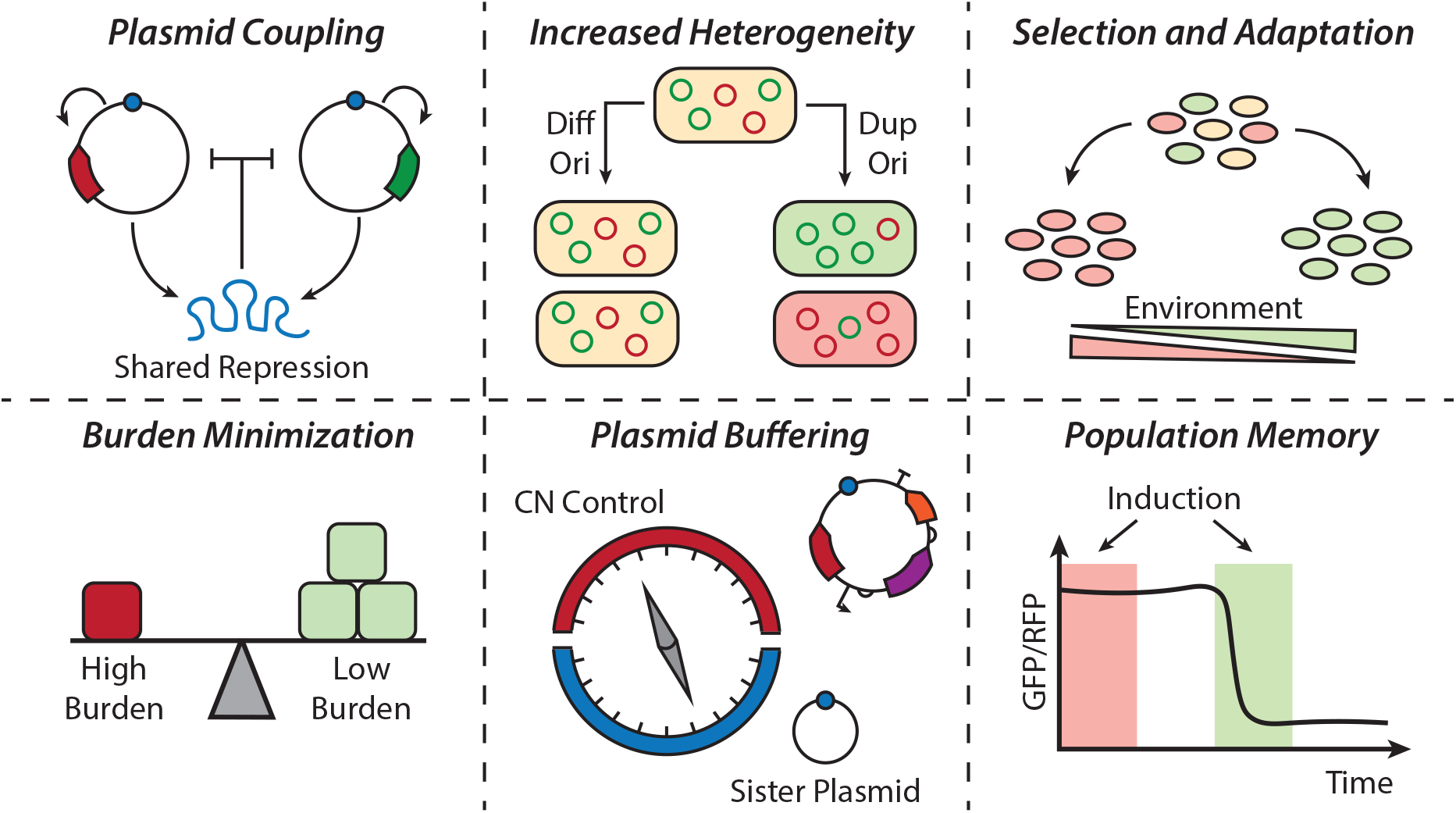
Overview of manuscript. Two plasmids are coupled via shared copy number regulation mechanisms. This coupling leads to increased phenotypic heterogeneity in a population. Different cells are selected for in varied environments due to fitness advantages. Systems with two incompatible plasmids allow for burden minimization of high cost plasmids. They also enable buffering of an essential plasmid of interest by a non-essential, minimal sister plasmid. Lastly, coupled plasmid systems can maintain population memory following selection.

## 1 Results

### Duplicate origin plasmid pairs generate population heterogeneity and enable environmental adaptation

Before engineering complex, two-plasmid systems with increased CN heterogeneity due to shared oris, we wanted to better understand the distribution of two distinct ColE1-type plasmids co-existing in a population of *E. coli*. To characterize heterogeneity in populations with duplicate ori constructs, we used two simple plasmids, one carrying a chloramphenicol resistance cassette (CmR) and expressing a red fluorescent protein (RFP) constituitively and one with a spectinomycin resistance cassette (SpecR) and expressing a green fluorescent protein (GFP) (Fig. 2A).

**Figure 2:**
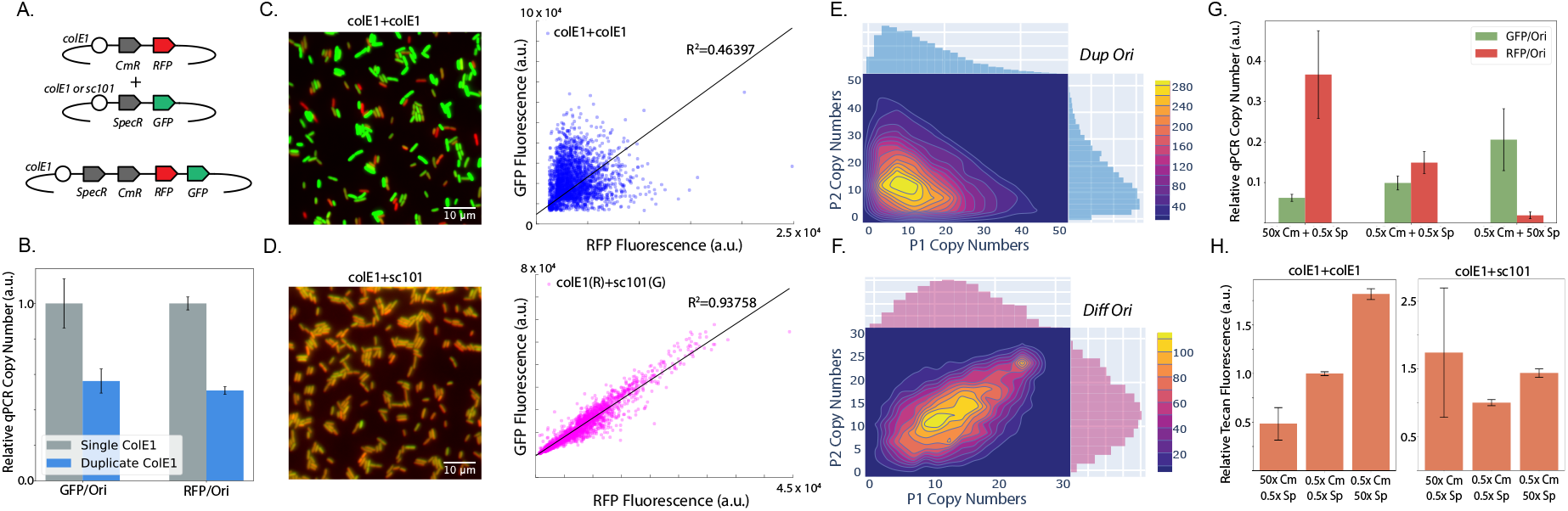
Two plasmid systems with redundant oris show increased heterogeneity and adaptability. (A) Diagram of plasmids used to look at CN behavior in different ori combinations. (B) Mean population CNs for a single ColE1 plasmid compared to two distinct ColE1 plasmids in the same cell as determined by qPCR. Error bars represent standard deviation of N=3 measurements. (C) Fluorescent micrograph of single cells with duplicate ColE1 ori plasmids and accompanying scatter plot showing GFP/RFP distribution for the population. (D) Fluorescent micrograph of single cells with different ori plasmids and accompanying scatter plot showing GFP/RFP distribution for the population. (E) Results from a model simulation showing CN distribution for incompatible plasmid pairs. (F) Results from a model simulation showing CN distribution for compatible plasmid pairs. (G) Bar chart showing relative plasmid CNs for the duplicate ori strain when grown in the presence of varying antibiotic concentrations. (H) Plate reader fluorescence for duplicate and different ori strains under different antibiotic concentrations.

To test the fundamental assumption that duplicate ori plasmids would share a coupled total CN similar to that of a single plasmid with the same ori, we performed quantitative real time PCR (qPCR) on strains containing either the two plasmids described above or a single plasmid carrying both antibiotic resistance and fluorescence markers (Fig. 2A). Relative qPCR CN of each fluorescent marker divided by the total ColE1 ori CN confirmed that in the duplicate ori case, the CN of each GFP or RFP plasmid was split roughly equally and each had about half of the total CN of a single ColE1 plasmid (Fig. 2B).

While qPCR provides a useful, population-level measurement of average CN, we wanted to better understand the underlying, single-cell CN distribution in two plasmid systems that either had the same oris or different oris with orthogonal regulation mechanisms. To do this, we compared two strains: one containing the GFP and RFP-expressing ColE1 plasmids described before (Dup ori) and one with an RFP-expressing ColE1 plasmid and a GFP-expressing sc101 plasmid (Diff_Ori) (Fig. 2A). To obtain single cell fluorescence measurements of the Dup Ori and Diff_Ori strains, we prepared agarose pad samples of each strain for fluorescence microscopy. In both the Dup Ori and Diff_Ori strain, we saw considerable variability in both GFP and RFP expression. For the Diff_Ori strain, GFP and RFP expression were well correlated (*R*^2^ = 0.9357) suggesting that variability in GFP and RFP expression was due to variability in the cell state that affected the CN of both plasmids similarly (Fig. 2D). Conversely, for the Dup Ori strain, GFP and RFP expression showed little to no correlation (*R*^2^ = 0.464), suggesting that the most significant cause of variability in this strain was the shared replication mechanism between the plasmids (Fig. 2C). Using a probabilistic plasmid replication and division model, we simulated growth of a population carrying two plasmids of either different or duplicate oris (Methods Supplemental Fig. 1). In the absence of selection, we saw that this simple theory recapitulated the divergent heterogeneity we found experimentally with duplicate ori plasmids (Fig. 2E,F).

Having shown that plasmid CNs in the Dup Ori strain were linked and heterogeneous, we wanted to see if the relative CN of the two plasmids could shift in response to different environmental conditions. To test whether variable antibiotic concentrations could bias the mean population CN, we cultured the Dup Ori strain in 50x the normal working concentration of chloramphenicol or spectinomycin. As expected, the population shifted its plasmid CN distribution towards the favorable plasmid (the one with the relevant resistance gene), as determined by qPCR (Fig. 2G).

Although absolute GFP and RFP fluorescence is likely affected by population health, the ratio of GFP to RFP is representative of the population’s CN distribution between the two plasmids. Plate reader measurements of bulk GFP/RFP fluorescence agreed with the qPCR results showing the population shift towards the more beneficial plasmid in each growth condition (Fig. 2H).

When looking at the response of the Diff_Ori strain to growth on different antibiotic concentrations, we did not see a significant difference in relative GFP and RFP fluorescence levels measured with a plate reader, indicating that this strain likely did not undergo significant changes in relative CN in response to different selection conditions (Fig. 2H). In summary, we showed that plasmids carrying the same ori type have coupled CNs leading to increased population heterogeneity and the ability to undergo fitness-based environmental selection and adaptation.

### Duplicate ori systems enable burden minimization and can increase plasmid stability due to buffering by non-essential plasmids

After showing fitness-based selection in response to antibiotic stress, we investigated whether duplicate ori strains could adapt to more biologically-relevant challenges, such as nutrient availability. It is well known that expression of metabolic pathway genes in *E. coli* is tightly regulated and dependent on metabolites in the surrounding environment. Furthermore, pathways for metabolizing nutrients can have complicated cost-benefit relationships, where cells must balance costly protein expression and enzyme activity with the benefit of increased nutrients for growth. For instance, Eames and Kortemme surprisingly found that for the lactose metabolizing operon of *E. coli*, lac permease activity is the most significant cost associated with operon expression and dictates cost/benefit trade-offs in lactose metabolism^17^. While native *E. coli* carbon source operon CNs are essentially constant (at one copy), modification of CN is presumably one mechanism cells could use to balance metabolic costs and benefits. We wanted to see if CN flexibility enabled by duplicate ori plasmid pairs would allow a population of *E. coli* to tune enzyme expression levels for improved growth on different carbon sources.

To this end, we created an *E. coli* strain that had the full arabinose operon (araOp) on a RFP-expressing ColE1-type plasmid and the full lactose operon (lacOp) on a GFP-expressing plasmid with the same ColE1 ori (Dup ori met). We transformed these plasmids into an *E. coli* host (JS006) that has the native genomic versions of these operons knocked out^18^.(Fig. 3A, Supplementary Fig. 2A). Naively, we expected the population to shift its CN distribution in favor of the essential carbon metabolizing plasmid while growing on that carbon source, for example increasing the CN of the araOp plasmid when grown on minimal media containing arabinose. However, when we grew this strain in batch culture in M9 media with different carbon sources we saw the opposite trend. Specifically, when grown on arabinose without antibiotic selection, we observed that GFP expression was significantly increased relative to growth on glucose or lactose and RFP expression was significantly decreased (Fig. 3B). When growing on arabinose, cells with fewer copies of the araOp likely had a fitness advantage over those with a higher plasmid operon CN. As a control, we created a strain that had the araOp on an sc101 ori plasmid and the lacOp on a ColE1 ori plasmid (Diff_ori_met), and thus should have no direct CN linkage. This effect was seen at a much smaller scale in the Diff_ori_met strain, but was just as pronounced in a duplicate ori strain with an MG1655 background that carries the genomic arabinose and lactose operons. Based on these results, we concluded that metabolic burden caused by induction of operon genes at high CN was the main driver of fitness-based adaptation in the batch culture, carbon metabolism experiments.

**Figure 3:**
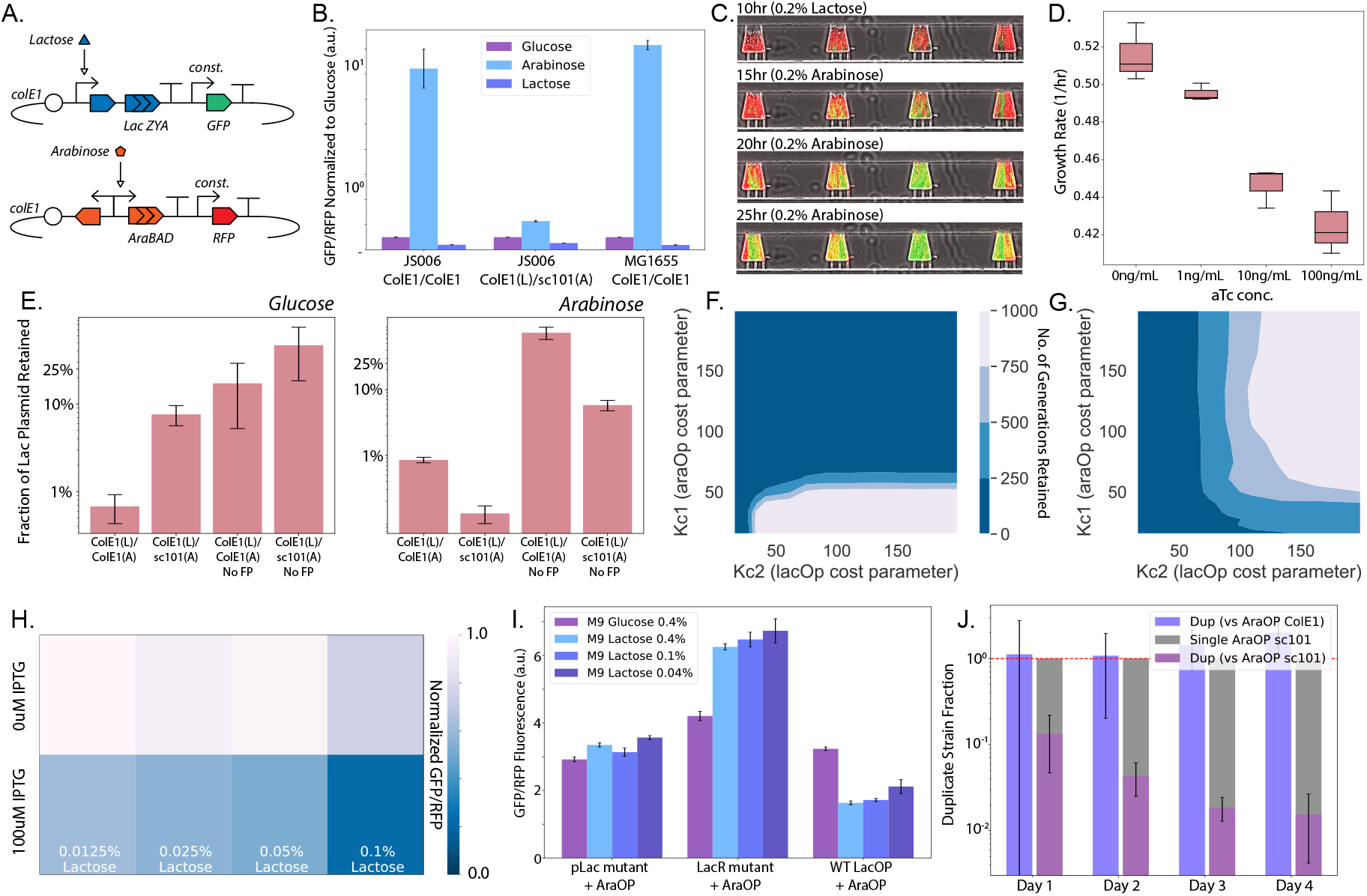
Duplicate origin plasmids allow burden-based adaptation to different nutrient environments and improve plasmid stability. (A) Circuit diagram for carbon source operon plasmids. GFP is expressed constitutively from the lactose operon-expressing plasmid and RFP is expressed constitutively from the arabinose operon-expressing plasmid. In the Diff_ori_met strain, the arabinose operon plasmid has the sc101 ori. (B) Plate reader fluorescence data for Dup ori met, Diff_ori_met, and Dup ori met MG1655 strains for growth on different carbon sources in M9 minimal media. Error bars represent standard deviation of three separate cultures. (C) Representative fluorescence micrographs for the Dup ori met strain grown in a microfluidic device. Culture media carbon source was switched from lactose to arabinose at t=10.5 hours. (D) Growth rate averages for ipUC-Ara strain whose CN is directly induced with increasing aTC concentration. (E) Plasmid retention data for Dup ori met and Diff_ori_met strains grown on different carbon sources in M9 minimal media. Error bars represent standard deviation of three separate cultures. (F) Simulation of lactose plasmid retention when grown in arabinose for duplicate ori plasmids. Colormap corresponds to number of generations with plasmid still retained. (G) Same simulation for different ori plasmids. (H) Heatmap showing plate reader fluorescence data for the Dup ori met strain for different concentrations of lactose and IPTG in M9 media. Each square represents the mean value of three separate cultures. (I) Bar chart showing plate reader fluorescence data for different versions of the Dup ori met strain with mutated lac operons for different concentrations of lactose. Error bars represent standard deviation of three separate measurements (J) Proportion of Dup ori met strain in population when cultured over time in competition with single plasmid arabinose operon strain.

To further investigate CN adaptation to carbon source type, we continuously cultured the Dup ori met strain in a microfluidic device^19^ that allowed us to vary the composition of growth media throughout the experiment (Fig. 3C). Cells were first cultured in M9 glucose, followed by media switches to M9 lactose and then arabinose. No antibiotics were included in this experiment. When grown on lactose minimal media, the cell traps in the device consisted primarily of strongly-expressing RFP, weakly-expressing GFP cells indicating higher araOp CN relative to lacOp CN. Notably, many cell traps also had smaller sub-populations of cells with stronger GFP expression than RFP expression indicating higher lac operon CN. In response to a media switch to M9 arabinose, growth halted in all cell traps, but about 15% of 40 analyzed cell traps were able to resume growth. In the traps that resumed growth. the strongly-GFP expressing subpopulations rapidly took over the cell trap to become the dominant phenotype (Supplement Video 1). These results supported our interpretation of the batch culture results that population CN heterogeneity promoted adaptation to different nutrient conditions.

Each operon is naturally found in the genome at one copy and therefore is likely expressed optimally at low CN. We hypothesized that cells with higher CN of the essential plasmid had considerable burden when the operon was turned “on” as compared to cells with lower CN of the essential plasmid and a high CN of the non-essential and transcriptionally “off” plasmid. For example, when the Dup ori met strain is grown in arabinose, it requires at least one copy of the araOp plasmid to survive, but selects for cells with high lacOp plasmid CN. Since the lacOp plasmid genes are turned off when lactose is not present, this plasmid acts as a buffer to drive down the CN of the more burdensome araOp plasmid. We call this feature of coupled plasmid systems *plasmid buffering*, and we are the first group to our knowledge to describe it.

To confirm our hypothesis that maintaining low CN of an essential araOp plasmid optimizes growth rate in arabinose minimal media, we built a version of our ColE1 araOp plasmid with direct aTc-inducible CN (Supplementary Fig. 2B) based on a recent publication^20^. When transformed into *E. coli* MgG1655 Z1, induction of CN with aTc led to a dose-dependent decrease in growth rate, suggesting that cells with a higher CN experience higher burden (Fig. 3D). These results further suggest that the Dup ori met strain used plasmid buffering as a strategy to balance the cost and benefit of two carbon-metabolizing plasmids in different environments.

Systems with duplicate oris are always avoided in synthetic biology due to more rapid loss of one plasmid from the population. However, our previous results with plasmid buffering suggested that there are certain conditions where plasmids with the same ori may exhibit increased stability compared to plasmids with different oris. Specifically, we hypothesized that, when grown on arabinose-only media, the Dup ori met strain would retain the lacOp plasmid for longer than the Diff_ori_met strain. Remarkably, we saw that the fitness advantage conferred by plasmid buffering can stabilize non-essential plasmids in duplicate ori systems without the need for antibiotics. We found that when grown in M9 glucose, the Dup ori met strain lost the lacOp plasmid faster than the Diff_ori_met strain, but when the strains were grown on M9 arabinose, Dup ori met retained the lacOp plasmid for longer than the Diff_ori_met strain(Fig. 3E). Contradicting conventional wisdom regarding incompatible plasmids, in this situation, the non-essential buffer plasmid is protected from loss by indirectly reducing the burden of the essential plasmid. When we repeated this plasmid loss experiment on M9 lactose, we did not see increased stability in the Dup ori met strain compared to the Diff_ori_met strain (Supplementary Fig. 2D). We hypothesize that this is because the lacOp plasmid is not as burdensome as the araOp plasmid at high CN.

After demonstrating the general ability of our Dup ori met strain to buffer the burden of highly expressed metabolic operons, we tested the ability of duplicate ori systems to optimize their CN distribution in response to a gradient of resource availability. To do this, we cultured the strain in varied concentrations of lactose from 0.0125% to 0.1% (Fig. 3H). Our results indicate a decrease in the proportion of lacOp plasmid as lactose availability increased, suggesting reduced need for multiple copies of the lacOp plasmid. To separate the potential effects of metabolic need from lactose inducer burden, we also cultured the strain in the same lactose gradient but with 100uM IPTG, which ensures lacOp genes are maximally expressed. The gradient CN adaptation response to lactose availability remained present with IPTG and showed an overall reduction in GFP/RFP due to full expression of the operon. When we repeated this experiment with the Diff_ori_met control strain, we did not see significant differences in GFP/RFP for different lactose concentration in the presence or absence of IPTG (Supplementary Fig. 2C).

In synthetic biology applications, plasmid constructs are usually not finely tuned to changing nutrient environments. Often it is hard to predict whether the expression level set by promoter choice is appropriate. In our WT version of the Dup ori met strain, lacOp expression was generally too costly. To investigate whether weakened versions of our metabolic constructs would optimize to a different CN point, we created two mutant versions of the lacOp plasmid, a pLac mutant known to decrease expression and a LacI mutant that also reduces operon expression^21^. These mutant lacOp ColE1 plasmids were co-transformed with a ColE1 ori araOp plasmid and the resulting strains were grown in various concentrations of lactose (Fig. 3H). Intuitively, both mutants with reduced operon expression had increased GFP/RFP, reflecting increased need for multiple copies of the weaker lacOp plasmid as compared to cells carrying the original plasmid. The LacI enhanced repression mutant also showed a gradient need for the lacOp that again increased with lowered lactose concentrations.

In summary, we have demonstrated the ability of duplicate ori strains to optimize the CN distribution between two carbon metabolizing constructs based on their expression strength and environmental growth condition. We also showed the benefits of plasmid buffering in reducing plasmid loss and improving strain fitness.

### Minimal sister plasmid provides copy number flexibility and evidence of population memory

Following the discovery of plasmid buffering, we wanted to examine whether we could use it as a tool to characterize the cost-benefit balance of an arbitrary synthetic gene circuit as a function of environment. To realize this, we introduced the idea of a “sister plasmid” that shares the same ori as an essential plasmid, but only contains that ori and a selection marker to minimize its burden on the cell. We hypothesized that sister plasmids could impart CN flexibility and thus promote sensitive adaptation to environment and also probe the environmentally determined cost-benefit function of an essential plasmid.

To test this idea, we co-transformed a ColE1 ori sister plasmid with a ColE1 plasmid carrying an AHL-inducible kanamycin resistance cassette (KanR) and constitutive GFP to make strain Dup ori kan (Fig. 4A). Using this setup, we directly varied the cost-benefit relationship of the plasmid in two directions by inducing different levels of KanR expression with AHL and also manipulating kanamycin concentrations in the culture media. In a plate reader experiment, we grew Dup ori kan in LB media containing 1x to 15x kanamycin working concentration and also 0nM to 100nM AHL. Convincingly, we saw that as kanamycin concentrations increased in the media, the KanR plasmid’s CN went up as represented by increased GFP/OD. Conversely, as expression of KanR was increased via AHL, the CN went down to compensate for burden due to excess KanR expression (Fig. 4B). After passaging these cells into 1X kanamycin media, we incubated them in the reverse conditions of the initial selection. For example, cells previously incubated in 15x kanamycin and 0nM AHL were now cultured in 1x kanamycin and 100nM AHL. The resulting GFP/OD measurements matched expected CN adaptation, showing reversibility and secondary adaptation of Dup ori kan to a new environmental set point. When this same experiment was performed on a control strain with a minimal sc101 ori sister plasmid (strain Diff ori kan), we saw much smaller changes in GFP/OD in response to different selection conditions (Supplementary Fig. 3).

**Figure 4:**
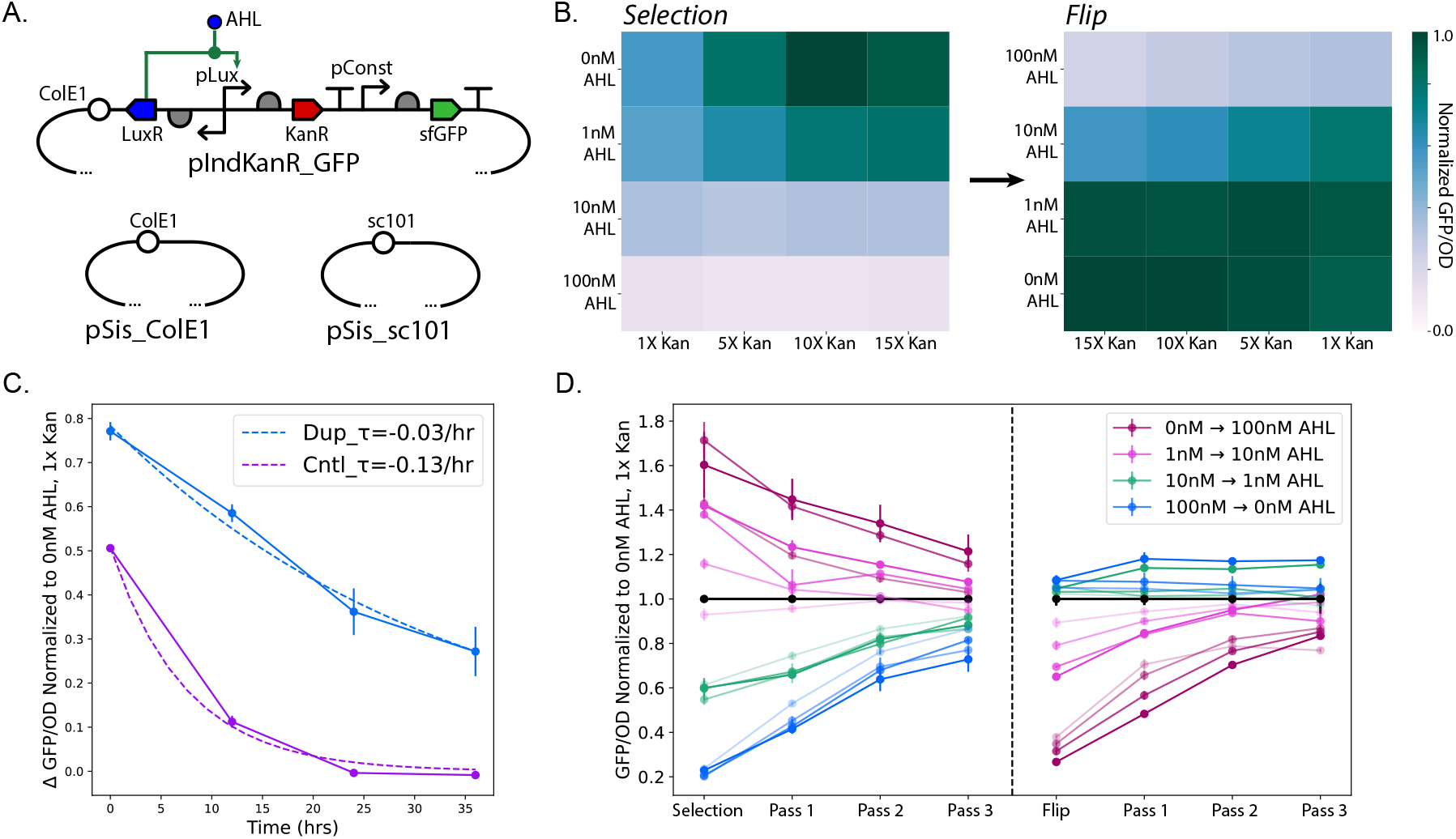
Incompatible “sister” plasmid buffering enables sensitive population adaptation and promotes long-term memory. (A) Circuit diagram of AHL-inducible KanR plasmid and accompanying minimal sister plasmids. (B) Heatmap showing plate reader fluorescence data for the Dup_ori_kan strain for different concentrations of AHL and kanamycin. Each square represents the mean of three separate cultures. Second heatmap shows fluorescence of cells following passage into “flipped” inducer conditions. (C) Representative plot of fluorescence decrease over time during passaging in non-selective media. Measurements of Dup_ori_kan and Diff_ori_kan strains are fit to exponential curve to estimate memory time constants. Error bars represent standard deviation of three separate cultures. (D) Plate reader fluorescence measurements for Dup_ori_kan in all inducer conditions over time, including non-selective passaging and subsequent flipped induction. Line opacity represents environmental kanamycin concentration with highest opacity corresponding to 15x Kan.

Based on these new results, we went back to improve our Dup ori met strain’s growth on arabinose using a minimal sister plasmid. We replaced the lacOp plasmid in that strain with a minimal, ColE1 sister plasmid so that the strain could only grown on arabinose, but could still tune araOp copy number. To see if this new duplicate ori strain had a fitness advantage when compared to a single plasmid strain (containing only the araOp plasmid on either an sc101 or Cole1 backbone), we competed the two in co-culture on arabinose minimal media. By plating the culture on spectinomycin plates versus antibiotic-free plates, we quantified the proportion of the population that was the new Dup ori ara sis strain over time. Over the course of 4 days, we saw that this Dup ori ara sis strain out-competed a single ColE1 araOp plasmid strain while it had a moderate fitness disadvantage compared to a single sc101 araOp plasmid strain with lower CN than ColE1 (Fig. 3I).

We then hypothesized that the use of sister plasmid buffering not only enabled reversible adaptation of plasmid CN to environmental need, but could also generate population memory of CN state. In the inducible KanR experiment described previously, GFP/OD was also measured over three non-selective passaging time points after each selection phase. Rather than immediately losing CN shifts once selection is removed, the population retained memory of the selection event and returned to baseline gradually over many generations (Fig. 4D). The time scale of population memory for Dup ori kan (0.03/hr) was considerably longer than in a distinct ori control strain (0.13/hr) where some CN adjustment was seen but was then rapidly lost upon removal of selection (Fig. 4C).

In summary, the use of a minimal sister plasmid with a duplicate origin enables the costbenefit characterization of synthetic constructs in different environments. In addition, plasmid coupling creates population memory of CN state without the need for additional circuit components.

### Redundant ori plasmid systems promote generational population memory in response to engineered cell lysis in microfluidic culture

To further investigate the temporal dynamics of population memory in continuous culture, we used a duplicate ori strain carrying two inducible lysis constructs on ColE1 plasmids (Dup ori lys). Each plasmid rapidly triggers cell death via the E lysis gene from phage *ϕ*X174 in response to a small molecule inducer. This model system creates easily tunable bidirectional selection as well as short time scales for adaptation due to the all-or-none nature of lysis-based selection. Together these attributes make plasmid lysis circuits optimal to investigate CN dynamics and memory in continuous culture. The first plasmid codes for AHL-inducible lysis and constitutive GFP, while the second encodes arabinose-inducible lysis and constitutive RFP (Fig. 5A). After confirming successful CN adaptation of Dup ori lys against lysis pressure in plate reader experiments (Supplementary Fig. 4A), we then cultured the strain in a previously described microfluidic chip along a gradient of arabinose and AHL concentrations (Fig.5B and Supplementary Fig. 5C). In this device, small bacterial populations grow as monolayers in cell traps of different sizes^22^. The device has two inlet media sources connected to a set of branching fluidic channels that create a concentration gradient across the different cell traps of the device, allowing us to vary arabinose and AHL concentrations on a single microfluidic chip. When cultured in this system, the Dup ori lys strain adapted similarly to batch culture, with high GFP-expressing, pAHL Lyse plasmid dominant populations taking over cell traps for high arabinose concentrations and high RFP-expressing, pAra Lyse plasmid dominant populations taking over cell traps for high AHL concentrations. Resulting GFP and RFP fluorescence over a 24 hour run are summarized in Fig. 5B for inducer concentrations up to 100nM AHL or 0.017% Arabinose. When we repeated this experiment with a control strain (Diff_ori_lys) that had the pAra lyse plasmid on an sc101 ori, we saw virtually no CN shifting to different arabinose and AHL conditions, further suggesting that ori redundancy enabled adaptation (Supplementary Fig. 4C and 5A)

**Figure 5:**
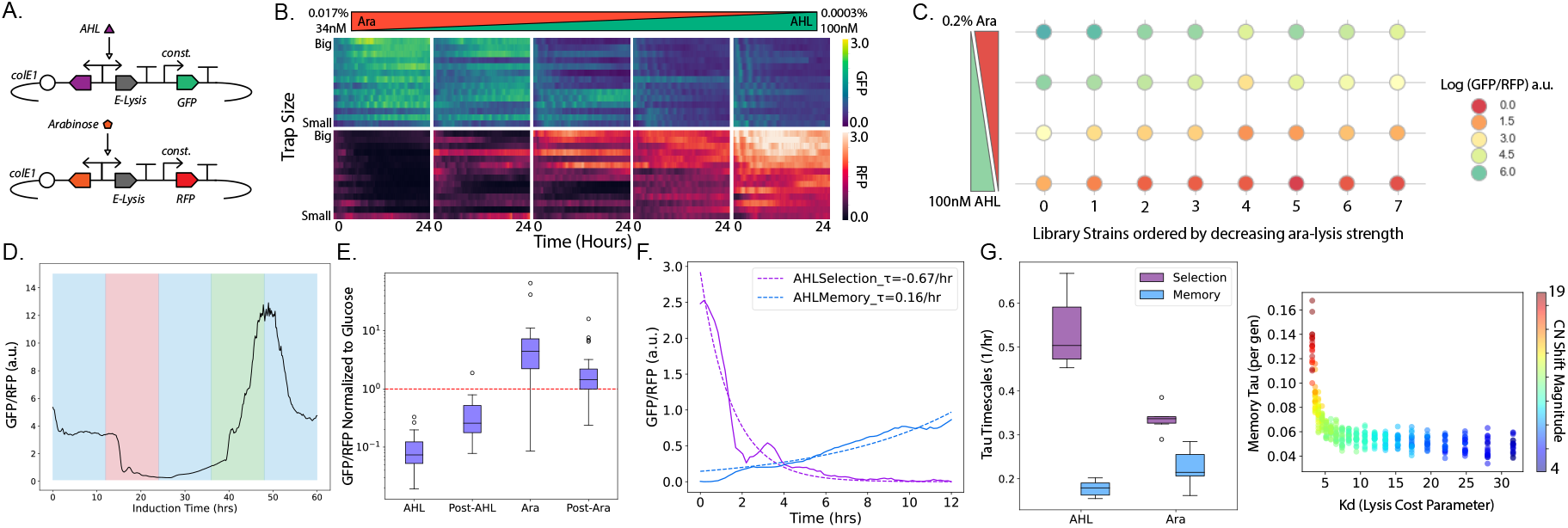
Inducible cell lysis triggers population CN shifting in duplicate ori systems. (A) Circuit diagram for inducible lysis plasmids. GFP is expressed constitutively from the AHL inducible lysis plasmid and RFP is expressed constitutively from the arabinose inducible lysis plasmid. In the Diff_ori_lys strain, the arabinose inducible lysis plasmid has the sc101 ori. (B) Heatmap showing mean fluorescence over time for the Dup_ori_lys strain in a microfluidic device with different inducer concentrations and trap sizes. (C) Plate reader fluorescence of Dup_ori_lys strain library with variant arabinose lysis strength across four inducer conditions. (D) Mean fluorescence over time for Dup_ori_lys strain in a microfluidic device with five induction windows: 0.2% Glucose (non-selective), 100nM AHL (selective), Glucose, 0.02% Arabinose (selective), and Glucose. (E) Mean fluorescence of Dup_ori_lys strain at the end of each induction window in previous panel. Mean and standard deviation of 24 individual traps are represented. (F) Representative mean fluorescence plot over time of one row of four traps during AHL selection and memory phases. Curves are fit to exponential function to estimate memory time constants. (G) Mean memory time constants for all 24 traps during each phase of microfluidic experiment. (H) Simulated relationship between memory time scale, CN shift magnitude, and lysis strength after 100 generations of selective growth and during 100 non-selective generations.

We hypothesized that due to rapid selection in this system, CN adaptation would sensitively respond to the lysis expression strength of each plasmid. In particular, we had found that the original Dup ori lys strain had significant leaky expression of the lysis gene, most likely due to expression from the arabinose promoter even when uninduced (Supplementary Fig. 4B). To investigate the sensitivity of CN adaptation to lysis strength we developed a library of pAra Lyse plasmids, each with different lysis gene expression strengths. This library was generated through site directed mutagenesis (SDM) of the ribosome binding site (RBS) before the E lysis gene on the original pAra Lyse plasmid. Each library plasmid’s lysis strength was assessed in response to induction with 0.02% arabinose as described in the methods (Supplementary Fig. 6A). These library plasmids were then co-transformed with the original pAHL Lyse plasmid and grown in various inducer concentrations in batch culture. A clear relationship between lysis expression strength and final GFP/RFP ratio was found in the 8 strains tested, with increasing arabinose lysis strength corresponding to a increase in GFP/RFP and therefore pAHL Lyse CN across all inducer conditions (Fig. 5C and Supplementary Fig. 6B).

After confirming the rapid and fine-tuned adaptation of Dup ori lys above, we characterized the strain’s memory dynamics using time-lapse microscopy. Specifically, we grew our strain along with Diff_ori_lys within a multi-strain microfluidic chip^19^. In 12 hour windows we cultured the populations first in LB media supplemented with glucose followed by LB with 100nM AHL, LB with glucose, LB with 0.02% arabinose, and LB with glucose again (Fig. 5D and Supplementary Fig. 4D and 4E). Antibiotics were used throughout the experiment to ensure at least one copy of each plasmid was always maintained. The Dup ori lys strain responded to both selection windows showing reversible adaptation. Again, we saw that following non-selective growth for 12 hrs even in high turnover continuous culture, populations still retained CN memory of the prior selection event, as measured by GFP/RFP levels (Fig. 5E). Following the first AHL induction, population memory was especially striking with the relative time scale of return to baseline being over four times slower than that of the selection phase (Fig. 5F). While the difference between selection and memory time scales for the arabinose induction phase was less substantial, this is likely due to the population being biased toward higher CN of the RFP before switching to arabinose and therefore undergoing slowed selection towards the pAHL Lyse GFP plasmid (Fig. 5G). The magnitude of GFP/RFP following arabinose selection still suggests memory is retained when compared to the initial glucose condition.

In summary, duplicate ori plasmids enable fast adaptation to engineered lysis burden in continuous culture experiments. The CN balance of these strains responds sensitively to the relative lysis gene expression strength of the plasmids and environmental conditions. By visualizing these populations over many generations in microfluidics, we could clearly demonstrate population CN memory following a selection event while still maintaining the ability to reversibly adapt. Overall this suggests the fundamental capability of plasmid coupling in producing population state memory over generational time scales without the need for added circuitry.

## Discussion

Synthetic biology as a field strives for tight control of cellular phenotypes. Gene circuits are finely tuned for deployment, and any variability is seen as a negative feature. In the natural world, however, populations will often exploit noise and heterogeneity in the face of unpredictable environments to improve survival. In this work we took inspiration from natural systems to create heterogeneous populations capable of environmental adaptation, burden minimization, and population memory. We used coupling between two plasmids of the same origin type to allow reversible copy number flexibility within a microbial population.

Previous work has shown that the expression level of metabolic proteins evolves towards an optimal level when cultured in a specific nutrient environment over time^23^. Here, we show optimization of protein expression strength based on environment by strains carrying duplicate origin plasmids that enable copy number adjustment rather than mutation to improve fitness. We introduced the concept of plasmid buffering where one essential plasmid driving population fitness is given a tunable copy number through the presence of a secondary non-essential plasmid with the same origin. Plasmid buffering in such systems allows for copy number adjustment in response to different growth environments. A recent study by Yang et al. created a synthetic circuit for eliminating gene dosage variation in individual mammalian cells^24^. Their method buffers plasmid CN variability in mammalian cells to reduce heterogeneity while we use plasmid CN flexibility to buffer circuit burden. Comparing these studies points out how copy number flexibility can be desirable sometimes, but copy number heterogeneity can be reduced if needed.

Recent work by the Ellis lab and others on both burden reporting and burden-responsive feedback control of synthetic circuits, highlights the importance of considering the effects of synthetic circuit burden on cellular fitness^25, 26^. Our study shows how plasmids with overlapping replication mechanisms can allow a cell to minimize burden caused by protein expression that is mismatched with the culture environment. In addition, we show that not only can plasmid buffering minimize burden, but it can also increase the stability of a non-essential plasmid by giving it an indirect benefit to the cell. Research on improving genetic stability of constructs has shown the effectiveness of reducing host mutational ability, overlapping sequences of essential and non-essential genes, and recoding translation among many other strategies^27^. Here we are able to enhance genetic stability specifically in duplicate origin systems. despite conventional wisdom regarding incompatible plasmids.

Plasmid incompatibility has been studied since the early 1970’s and there are multiple, published models to understand the dynamics of plasmid loss between plasmids with shared replication mechanisms^28–30^. Specifically, the effects of different methods for choosing which plasmid to replicate and how to partition these plasmids to daughter cells has been investigated mathematically^31^. However, there is a lack of mathematical models that consider the cost and benefit associated with plasmid encoded functions and how these costs and benefits can change with environment. Here we show that a basic probabilistic model that uses Hill-like functions to simulate plasmid cost and benefit can recapitulate many features of incompatible plasmids seen experimentally. In the future the model could be improved by trying to get more precise parameters for plasmid replication rates, cell growth rates, and cost/benefit function parameters. We could also include a more complex framework for host-circuit burden coupling, like some recently proposed approaches^32^.

While we are the first group to our knowledge to document adaptation due to duplicate ori plasmid pairs, a recent study by Tomanek et al. investigated a similar phenomenon of gene copy number adaptation^6^. Specifically, they describe a mechanism called amplification-mediated gene expression tuning (AMGET). While we showed that plasmid CN heterogeneity enables adaptation to variable environments, they “show that gene duplications and amplifications enable adaptation to fluctuating environments by rapidly generating copy-number and, therefore, expression-level polymorphisms.” The core idea of stochastic, copy number heterogeneity promoting population adaptation is shared between their study and ours. However, they describe different mechanisms for generating copy number variation. While we exploit the cell’s inability to distinguish plasmids with the same origin, Tomanek et al. studied how gene duplication and amplification arising from homologous recombination between sister chromosomes leads to CN variation. This mechanism of generating copy number occurs naturally in cells without synthetic gene circuits, which begs the question: are plasmid systems with redundant replication mechanisms also utilized in nature to cope with varying environments?

Lastly, we demonstrate memory in cell populations containing duplicate origin plasmids following environmental selection. Researchers have created population-state memory in synthetic systems through the use of bistable genetic circuits, and more recently the development of a methylation-based epigenetic system within *E. coli*^33^. Here, we develop generational memory through plasmid inheritance of population copy number distribution. Studies on history-dependence of microbial populations have cited chromatin state, protein inheritance, metabolic history, and strong heterogeneity as potential mechanisms for memory of past environmental exposures^34^. In this work, we took inspiration from these natural mechanisms of population adaptation to demonstrate the broad use of duplicate origin plasmids as a tool to improve performance of synthetic microbial populations.

## 2 Materials and Methods

### Strains and plasmids

*E. coli* MG1655 (NCBI U00096.3) was used as the host strain for the majority of experiments except where noted. For experiments involving the lactose and arabinose operon strains, *E. coli* strain JS006?,^18^ was used .This strain is derived from the *E. coli* Keio knockout collection parent strain, BW25113, and has the following relevant mutations: Δ(araD-araB)567,ΔlacZ4787(::rrnB-3), ΔlacI, ΔaraC. For experiments involving the inducible pUC copy number plasmids, the *E. coli* K-12 MG1655Z1 strain was used, which constituitively expresses lacI and tetR from the genome.

All plasmids were constructed by Gibson assembly using PCR amplified fragments of previously constructed plasmids from our group. The full *E. coli* lactose operon was amplified from the MG1655 strain genome. The full E. *coli* arabinose operon was also amplified from the MG1665 strain genome. The inducible lysis library was created using a mutagenic primer targeting the ribosome binding site in front of its lysis gene.

### Plate reader experiments for estimating average copy number in batch culture using fluorescence

For plate reader experiments assessing strain growth and bulk fluorescence, saturated overnight cultures of a given strain were diluted 1:100 into 200uL of fresh media and cultured in 96-well flat bottom plates at 37C with orbital shaking in a Tecan Infinite M200 Pro. Optical density (OD) and fluorescence measurements were taken every 5 minutes and the cultures were grown to saturation. GFP and RFP measurements were normalized by dividing by the culture OD at each time point. For use in comparative plots between conditions, final GFP/OD and RFP/OD were determined from when the OD maximized, or stopped increasing, a signal of early stationary phase.

### Agarose pad slide preparation for single-cell imaging and analysis

To image live *E. coli* cells following growth in relevant media conditions, overnight cultures were diluted 10 fold in sterile water before 2uL were pipetted onto a glass coverslip. Cells were then sandwiched by placing an agar pad (1.5% agarose) on top to create a flat focal plane. Agar pads were made the depth of one coverslip using the method described in ^35^. To analyze single cell fluorescence from images, a custom Matlab script was written that segmented single cells prior to recording their average fluorescent intensity. The segmentation workflow included image filtering, edge detection followed by object filling, and lastly individual cell classification and assignment using size and fluorescence thresholds.

### Quantitative PCR for determination of average copy number

For direct quantification of plasmid copy number, saturated cell cultures were analyzed by qPCR. Following growth in relevant media conditions, cells were incubated at 95C for 10 min prior to freezing at −20C. Lysed cultures were then diluted 100 fold for use as final templates. Each qPCR reaction consisted of 10uL of SYBR green master mix, 4uL water, 5uL diluted template, and 0.5uL of each primer. Primers were designed to amplify approx. 100bp regions within the GFP gene, RFP gene, plasmid origin, and the *E. coli* chromosome. Reactions were run in triplicate using MicroAmp Fast Optical 96-well plates within the Applied Biosystems Quant 3 machine for comparative CT measurements. Resulting measurements were normalized to chromosomal control reactions.

### Microfluidics and microscopy for carbon-source operon strains in multi-strain device

For time-lapse microscopy experiments with carbon-source operon strains, another previously developed device was used^19^. This device allows for observation of multiple strains within individual traps that are grown with the same inlet media source without cross-contamination. Carbon source experiments consisted of 24 hour induction periods with various supplemented M9 media.

### Serial passaging and plating experiment to estimate plasmid loss rate for carbon-source operon strains

Carbon metabolizing operon strains were grown overnight in 3mL of LB media with chloramphenicol and spectinomycin directly from glycerol stocks. The next day, the overnight culture was washed twice in M9 minimal media and used to inoculate new cultures at a 1:100 dilution in M9 media that was supplemented with Wolfe’s Vitamin Solution, MOPS for trace minerals, and either 0.2% of glucose, arabinose, or lactose (with no antibiotics). The cultures were grown for around 24 hours and passaged 1:100 into fresh, supplemented M9 minimal media that contained the same carbon source that the strain was grown in on the previous day. After 5 passages, samples of the different cultures were plated on LB agar plates containing either no antibiotics, chloramphenicol, spectinomycin, or both antibiotics to determine colony forming units (CFUs). Specifically, to determine CFUs, 3uL droplets of different serial dilutions of the culture were spotted on the agar plates and left to dry before culturing the agar plates at 37C overnight. For each culture, all of the different serial dilutions were plated in triplicate. The following day, CFUs in the culture were determined by counting the number of colonies that formed for the serial dilution that led to between 10 and 30 colonies on average. The number of CFUs in the undiluted culture was then determined by multiplying the number of counted colonies by the dilution factor. Finally, CFUs in antibiotic were divided by the total CFUs found on just LB to determine plasmid retention.

### Serial passaging and plating experiment to estimate population fraction of duplicate versus single carbon-source operon strains in competition

Duplicate origin carbon-source strains and single colE1 arabinose plasmid strains were grown separately overnight in supplemented M9 arabinose. They were then diluted 1:100 fold and co-cultured at an equal ratio in fresh M9 arabinose media with no antibiotics. Similar to the spot dilution method described for plasmid loss estimation, cultures were passaged every 24 hours and a sample was plated onto LB plates containing no antibiotic or spectinomycin. The next day colonies were counted to determine CFUs for all strains on LB or just the duplicate origin strain on LB-spectinomycin. Fraction of the population retaining the duplicate origin strains was then calculated each passaging day for 4 days.

### Serial passaging and memory quantification of inducible Kan strains in plate reader

For memory experiments with inducible Kan strains, overnight cultures were diluted 1:100 into selective media conditions containing AHL (0-100nM) and Kanamycin (1-15x) within a 96-well plate. After 12 hrs of growth, cells were diluted into 1x kanamycin, 0nM AHL media in a fresh 96-well plate. Three non-selective passages of 12 hrs each were carried out prior to incubation of cells in flipped selective media conditions. This selective phase exposed each culture to the opposite condition as the initial selective phase, for example cultures first incubated at 100nM AHL, 1x Kan were now exposed to 0nM AHL, 15x Kan. Following this secondary selective incubation, three more non-selective passages of 12 hrs each were performed. GFP/OD measurements were then plotted over time and each non-selective memory phase was fit to an exponential curve to determine the memory time constant (return rate) following GFP response.

### Microfluidics and microscopy for inducible lysis strains in concentration gradient device

For time-lapse microscopy experiments with the inducible cell lysis strains, a previously developed microfluidic device was used^?^. This device consists of separate arrays of small cell traps that are maintained at different inducer concentrations depending on the inducer concentrations in two upstream inlet channels of the device.

### Image analysis of inducible lysis strain timelapse microscopy experiments

To quantify population fluorescence from microfluidics experiments, we developed a FIJI macro. This macro created ROIs around each individual trap, thresholded fluorescence in each channel to select only healthy cells, merged these ROIs and measured mean intensity for each time slice. One value each for threshold minimum and maximum was set for an entire experiment to avoid non-cell background and also exclude unhealthy filamentous cells. Post processing of results also included background value subtraction for each trap and time slice.

### Microfluidics and microscopy for inducible lysis strains in multi-strain device

For time-lapse microscopy experiments with inducible lysis strains for memory investigation, the multistrain chip described previously was used again. Lysis memory experiments consisted of 12 hour induction periods with either non-selective LB-0.2% glucose media, 100nM AHL media, or 0.02% arabinose media. Duplicate ori and different ori strains were imaged during the same experimental run.

### Plasmid Copy Number Model

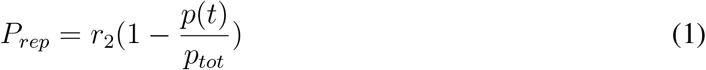

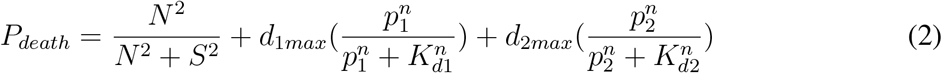

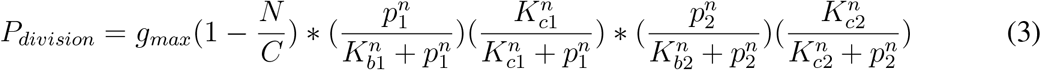

To study how the plasmid copy number distribution of a population changes over time, we created a stochastic model that takes into account the cost/benefit associated with a given plasmid. The model consists of cells containing two distinct plasmids that can have either the same or orthogonal mechanisms of replication and copy number maintenance. In each generation the following steps are carried out: 1) plasmids are replicated, 2) some members of the population die (due to plasmid encoded or effects of population saturation), 3) cells have the chance to divide based on plasmid-associated cost-benefit functions, and 4) for dividing cells, plasmids are partitioned to two daughter cells. We assume that the plasmid replication probability is given by Equation 1 where *r*_2_ is the maximum replication probability, *p*(*t*) is the current plasmid copy number and *p_tot_* is the maximum plasmid copy number. For “incompatible plasmids”, *p_tot_* is the same between plasmids whereas for compatible plasmids, each plasmid has its own maximum copy number. The number of plasmids that replicate in a given cycle is a random number from the binomial distribution with probability *P_rep_* and *p*(*t*) trials.

Equation 2 describes the probability for a given cell to die based off its current plasmid copy numbers. The first term represents death probability due to general saturation effects as the population size approaches the carrying capacity *C*. The parameter *S* represents the population size where the death probability due to culture saturation is half maximal. The two subsequent terms represent death probability due to lysis gene expression from plasmids *p*_1_ and *p*_2_ respectively. Death probability due to plasmid gene expression is assumed to follow the Hill Equation where *K_d_* represents the plasmid copy number that leads to half-maximal death probability and *d_max_* represents the maximum death probability.

Division probability is calculated in a similar manner to death probability as shown in Equation 3. In this equation, we assume logistic growth of the bacterial population where *g_max_* is the maximum division probability and *C* is the carrying capacity. Each plasmid *p*_1_ and *p*_2_ is assumed to have a potential benefit and cost that can increase or decrease the division probability. The Hill Equation is used to model plasmid cost-benefit effects where for each plasmid *K_b_* represents the plasmid copy number that leads to half-maximal benefit and *K_c_* represents the plasmid copy number leading to half-maximal cost. A cell divides if a number of one is randomly chosen from a binomial distribution with probability *P_division_* and one trial. If a cell is chosen to divide, its plasmids are partitioned to two daughter cells binomially.

## Abbreviations

Ori: Origin of replication
CN: Copy Number

## Acknowledgements

This work was supported by the National Institute of General Medical Sciences of the National Institutes of Health (grant no. R01GM069811). Andrew Lezia was supported in part by the National Science Foundation Graduate Research Fellowship Program (NSF GRFP)under Grant No. (DGE-2038238). Shalni Kumar was supported in part by the NSF GRFP under Grant No. (DGE-2038238). Shalni Kumar was also supported by the NIH-sponsored Quantitative Integrative Biology Training Grant (No. 5T32GM127235). The authors would like to thank Dr.’s Nick Csicsery and Ricky O’Laughlin for critically reading the manuscript and providing valuable feedback.

## Notes

### Competing Interest Statement

J.H. is a founder of GenCirq and Quantitative BioSciences, which focus on cancer therapeutics and agricultural synthetic biology, respectively. J.H. is a shareholder in both of these companies and is on their scientific advisory board.

