## Supplemental Figures for "Exploiting heterogeneity in coupled, two plasmid systems for dynamic population adaptation"

### Supplement

#### Supplementary Figure 1

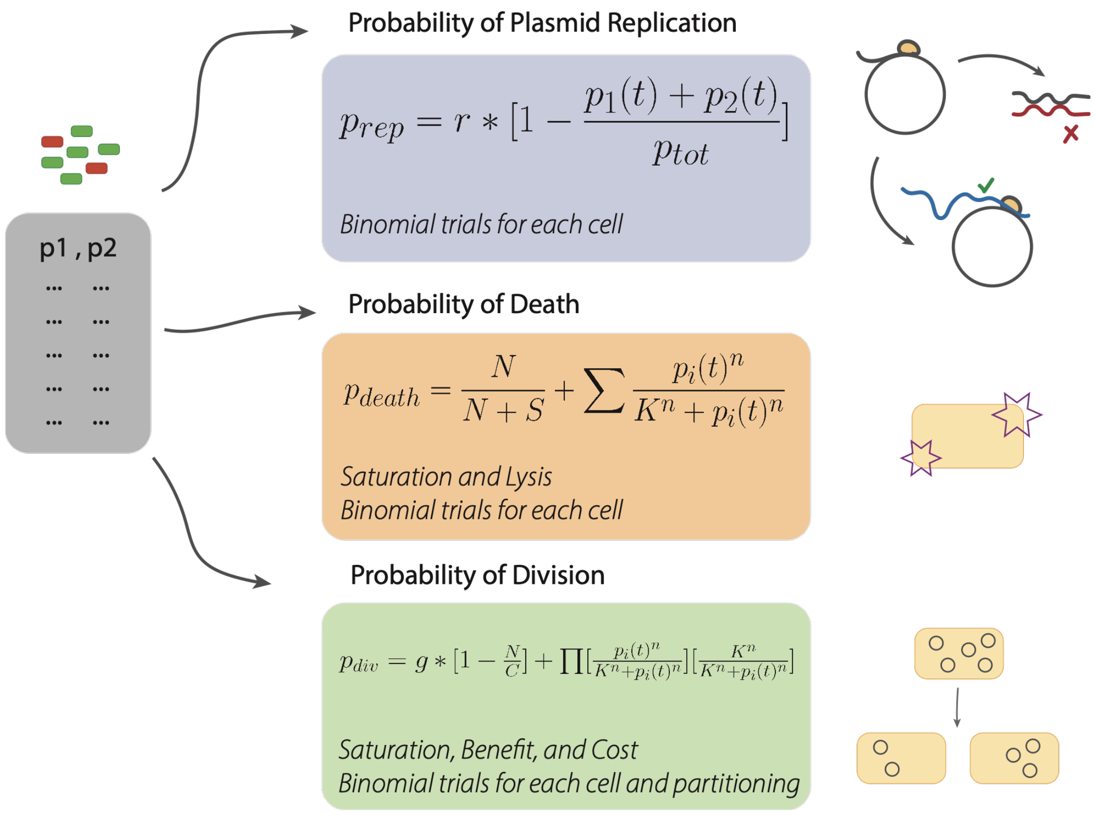


**Caption:** To study how the plasmid copy number distribution of a population changes over time, we created a stochastic model that considers the cost/benefit associated with a given plasmid. The model consists of cells containing two distinct plasmids that can have either the same or orthogonal mechanisms of replication and copy number maintenance. In each generation the following steps are carried out: 1) plasmids are replicated, 2) some members of the population die (due to plasmid encoded or effects of population saturation), 3) cells have the chance to divide based on plasmid-associated cost-benefit functions, and 4) for dividing cells, plasmids are partitioned to two daughter cells. See methods section for details on parameters.

#### Supplementary Figure 2

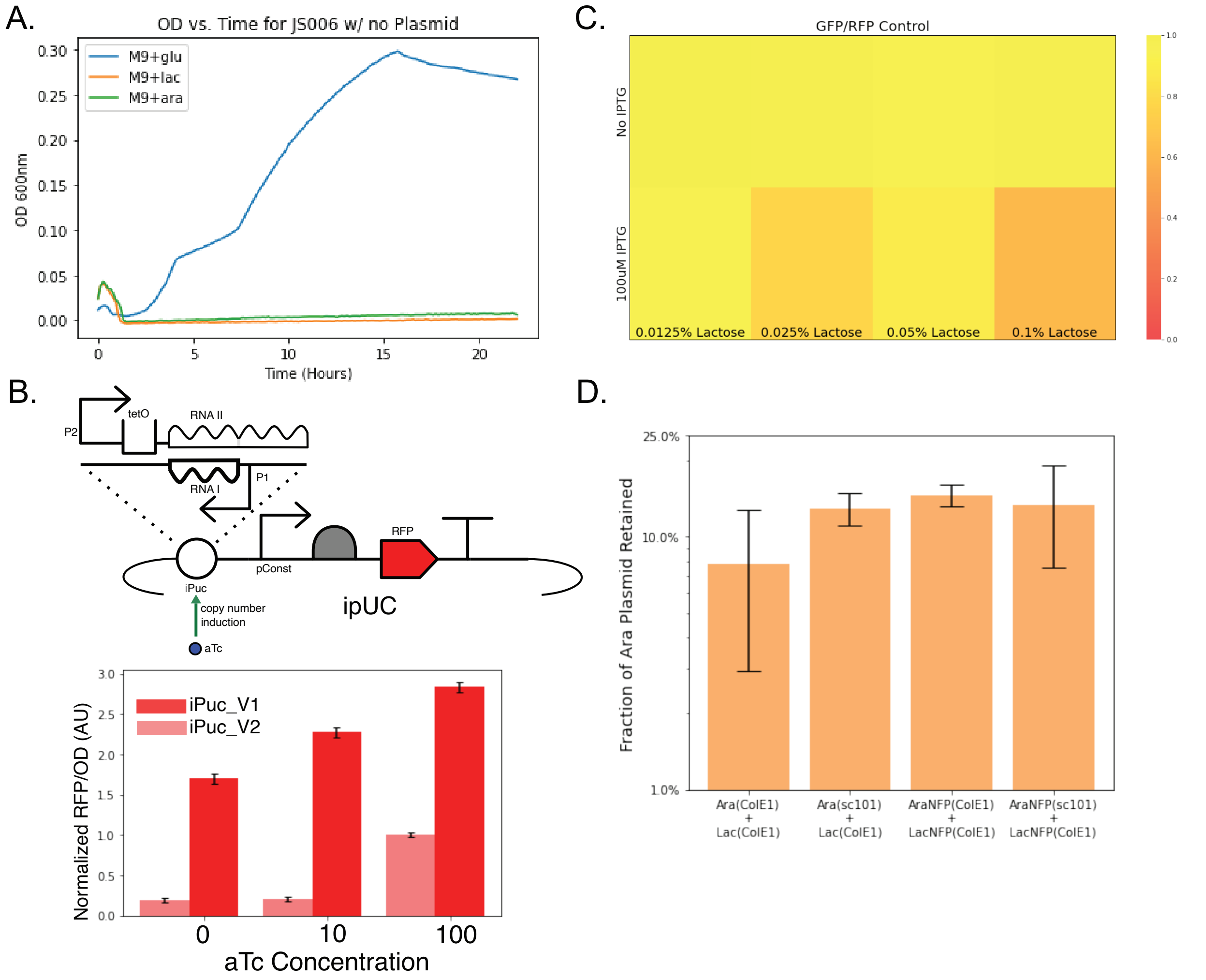


**Caption:** Supplementary material related to Figure 3. **(A)** Strain JS006 is unable to grow on minimal media with arabinose or lactose as the only carbon source unless operon genes are supplied on plasmids. **(B)** Proof of function for inducible copy number plasmid, Top: Circuit diagram for testing aTc-inducible pUC copy number. Bottom: Bar chart showing RFP fluorescence vs. aTc concentration for two different versions of the inducible pUC system with different tetO operator sequences. Error bars represent standard deviation of N=5 measurements. **(C)** Heatmap of GFP/RFP fluorescence values for the Diff_ori_met strain grown on different concentrations of lactose and IPTG. Comparison data for main Figure 3H. **(D)** Plasmid retention data for Dup_ori_met and Diff_ori_met strains grown on lactose minimal media. Comparison data for main Figure 3E.

#### Supplementary Figure 3

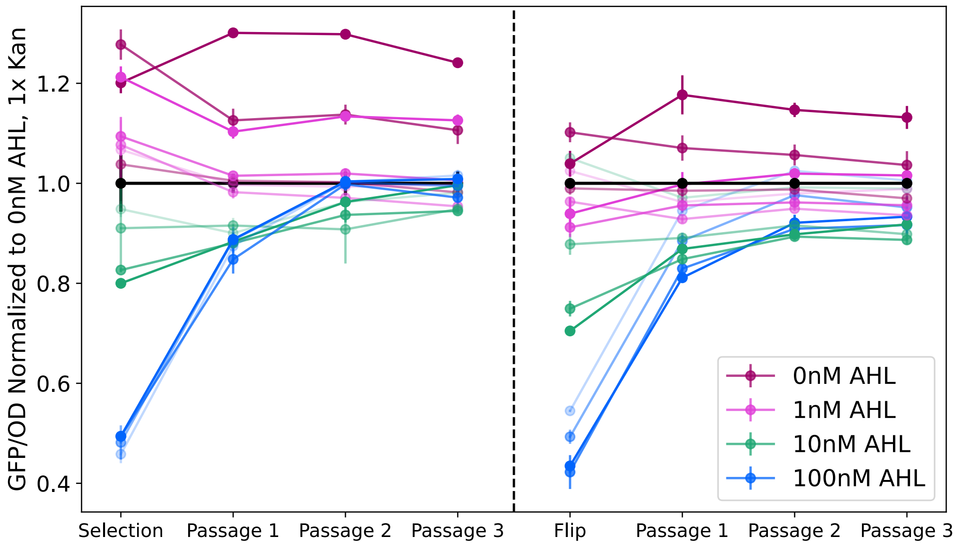


**Caption:** Supplementary material related to Figure 4. Plate reader fluorescence measurements for Dup ori kan in all inducer conditions over time during non-selective passaging with subsequent flipped induction. Comparison data for Figure 4D.

#### Supplementary Figure 4

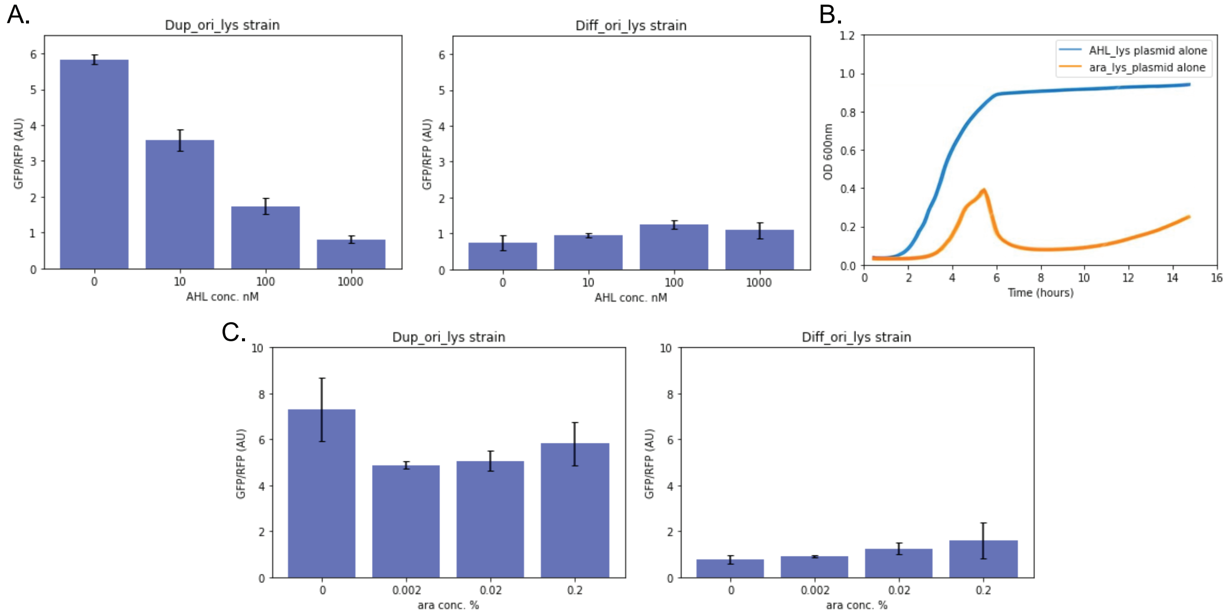


**Caption:** Supplementary material related to Figure 5. **(A)** Relative GFP/RFP fluorescence for the Dup_ori_lys and Diff_ori_lys strains grown in batch culture in different concentration of AHL. **(B)** Growth curves for strains containing either the pAHL_Lyse plasmid or the pAra_Lyse plasmid only. Even without inducer, the original pAra_Lyse plasmid causes cell death in LB as seen in the growth curve. **(C)** Relative GFP/RFP fluorescence for the Dup_ori_lys and Diff_ori_lys strains grown in batch culture in different concentration of arabinose.

#### Supplementary Figure 5

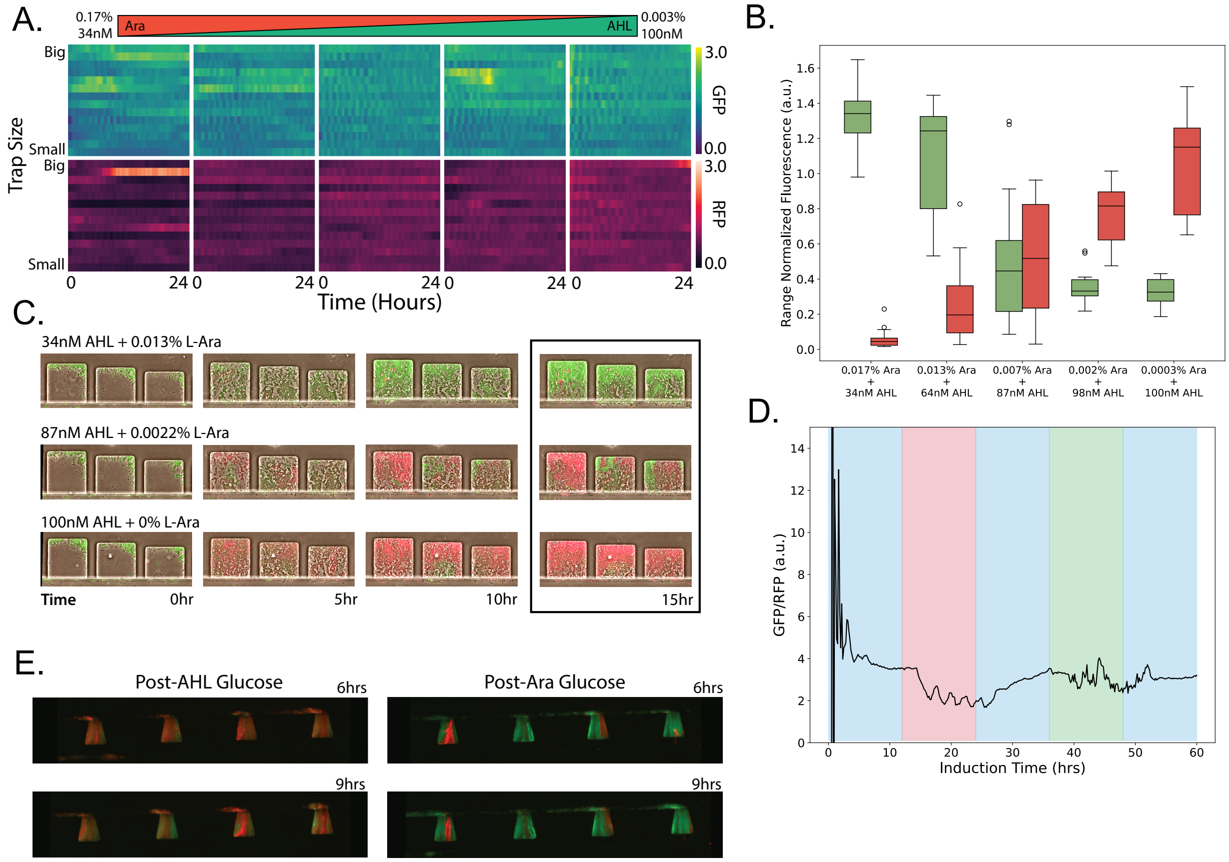


**Caption:** Supplementary material related to Figure 5. **(A)** Heatmap showing GFP and RFP fluorescence levels over time for the Diff_ori_lys strain grown in various concentration of AHL and arabinose in a microfluidic device. Comparison data for Figure 5B. **(B)** Boxplot showing final mean cell trap GFP/RFP fluorescence ratios for the Dup_ori_lys strain gradient microfluidic experiment. Associated with Figure 5B. **(C)**Representative microscope images from gradient microfluidic experiment with Dup_ori_lys strain. Images correspond to data shown in Figure 5B. **(D)** Mean fluorescence over time for Diff_ori_lys strain in a microfluidic device with five induction windows: 0.2% Glucose (non-selective), 100nM AHL (selective), Glucose, 0.02% Arabinose (selective), and Glucose. Comparison data for Figure 5D. **(E)** Representative fluorescence micrographs of the Dup_ori_lys strain grown in a microfluidic device over time in fluctuating environmental conditions. Microscope images correspond to data shown in Figure 5D-F. **(F)**

**Supplementary Figure 6**

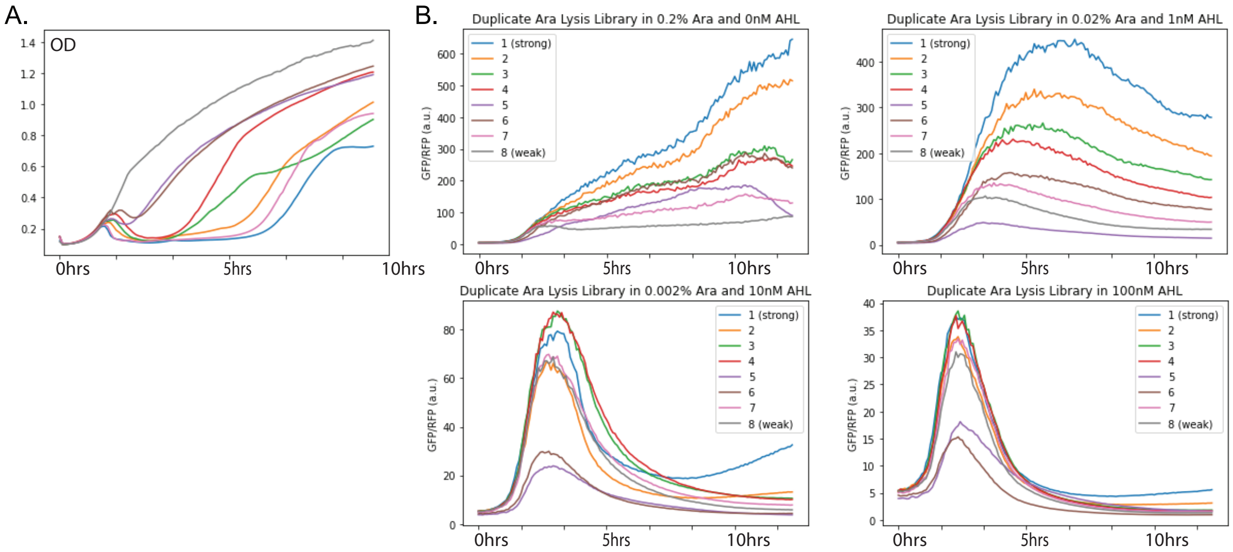


**Caption:** Supplementary material related to Figure 6. **(A)** Growth curves for 8 different strains grown in LB with 0.02% arabinose with various versions of the pAra_lyse plasmid. Each strain has a different RBS preceeding the lysis gene, E. Data related to Figure 5C **(B)** Plate reader fluorescence data over time for the 8 different Dup_ori_lys library strains grown in different AHL and arabinose concentrations. Data related to Figure 5C.

**Supplementary Table 1**

| Strain Name | Plasmids | Host Designation | Relevant Figure |
|---|---|---|---|
| Dup_ori | pSKAL001, pSKAL003 | MG1655 | 2 |
| Diff_ori | pSKAL002, pSKAL003 | MG1655 | 2 |
| Diff_ori_met | pSKAL006, pSKAL007 | JS006 | 3 |
| Dup_ori_met | pSKAL005, pSKAL006 | JS006 | 3 |
| Dup_ori_lys | pSKAL017, pSKAL019 | MG1655 | 5 |
| Diff_ori_lys | pSKAL017, pSKAL018 | MG1655 | 5 |
| Dup_ori_kan | pSKAL014, pSKAL015 | MG1655 | 4 |
| Diff_ori_kan | pSKAL014, pSKAL016 | MG1655 | 4 |
| Dup_ori_ara_sis | pSKAL005, pSKAL021 | JS006 | 3 |

**Caption:** Supplementary table detailing strains used in this study. Transformed plasmids are further described in supplementary table 2. JS006 host is a knockout for the lactose and arabinose genomic operons.

**Supplementary Table 2**

| Plasmid ID | Resistance | Origin | Fluorescent Protein | Circuit | Relevant Figure(s) |
|---|---|---|---|---|---|
| pSKAL001 | SpecR | ColE1 | GFP | N/A | 2 |
| pSKAL002 | SpecR | sc101 | GFP | N/A | 2 |
| pSKAL003 | CmR | ColE1 | RFP | N/A | 2 |
| pSKAL004 | CmR, SpecR | ColE1 | GFP, RFP | N/A | 2 |
| pSKAL005 | CmR | ColE1 | RFP | araBAD operon | 3 |
| pSKAL006 | SpecR | ColE1 | GFP | Lac operon | 3 |
| pSKAL007 | CmR | sc101 | RFP | araBAD operon | 3 |
| pSKAL008 | CmR | iPuc | RFP | araBAD operon | 3 |
| pSKAL009 | CmR | ColE1 | N/A | araBAD operon | 3 |
| pSKAL010 | CmR | sc101 | N/A | araBAD operon | 3 |
| pSKAL011 | SpecR | ColE1 | N/A | Lac operon | 3 |
| pSKAL012 | SpecR | ColE1 | GFP | Lac operon pLac mutant | 3 |
| pSKAL013 | SpecR | ColE1 | GFP | Lac operon LacI mutant | 3 |
| pSKAL014 | SpecR, (KanR) | ColE1 | GFP | AHL inducible KanR | 4 |
| pSKAL015 | CmR | ColE1 | N/A | N/A | 4 |
| pSKAL016 | CmR | sc101 | N/A | N/A | 4 |
| pSKAL017 | SpecR | ColE1 | GFP | AHL inducible lysis | 5 |
| pSKAL018 | CmR | sc101 | RFP | Arabinose inducible lysis | 5 |
| pSKAL019 | CmR | ColE1 | RFP | Arabinose inducible lysis | 5 |
| pSKAL020 | CmR | iPuc | RFP | N/A | Supplement |
| pSKAL021 | SpecR | ColE1 | N/A | N/A | 3 |

**Caption:** Supplementary table detailing all plasmids used in this study. Table includes annotations for plasmid antibiotic resistance, origin type, fluorescent proteins expressed, and any relevant circuits.

**Supplementary Table 3**

| **qPCR TABLE PRIMERS** | |
|---|---|
| GFP_fwd | GGGTGAAGGTGATGCTACAA |
| GFP_rev | GAACACCATAGGTCAGAGTAGTG |
| RFP_fwd | CACCCAGACCATGAGAATCAA |
| RFP_rev | TGGGTGTGGTTGATGAAGG |
| ColE1_fwd | CACGCTGTAGGTATCTCAGTTC |
| ColE1_rev | GGTTGGACTCAAGACGATAGTT |
| Chrom_fwd | GCGAGCGATCCAGAAGATCT |
| Chrom_rev | GGGTAAAGGATGCCACAGACA |

**Caption:** Supplementary table with primers used for qPCR in Figure 2. Each forward and reverse pair amplifies a small target region of ~100bp from plasmids within the sample of interest.
